# *Vibrio natriegens* is sensitive to its acidic fermentation products

**DOI:** 10.1101/2025.08.29.673078

**Authors:** Nicholas W. Haas, James B. McKinlay

## Abstract

*Vibrio natriegens* is an emerging bacterial platform for a range of biotechnological applications due to its rapid growth and ease of genetic manipulation. Whereas much has been learned about *V. natriegens* aerobic physiology, comparatively little is known about its anaerobic fermentative physiology, despite the relevance to many industrial conditions. We compared the metabolic parameters of *V. natriegens* versus another biotechnologically relevant bacterium *Escherichia coli* under fermentative conditions. Both species excreted a similar array of fermentation products but *V. natriegens* consumed less glucose and had a lower product titer. *V. natriegens* also exhibited rapid death, reaching extinction within 12 hours after the growth phase, 3 days sooner than *E. coli*. Rapid *V. natriegens* death was avoided, and glucose consumption and product titers improved, by increasing the buffering capacity of the growth medium, indicating that *V. natriegens* is comparatively sensitive to its organic acid fermentation products. Indeed, the *V. natriegens* genome lacks nearly all the acid resistance genes that have been characterized in *E. coli*. Our findings thus highlight an acid sensitivity that will need to be considered when designing fermentative applications of *V. natriegens*.

**IMPORTANCE:** Bioprocessing, the biological conversion of renewable resources into value-added chemicals, is poised to meet an increasing demand for sustainable alternatives to petroleum-based products. Many examples of bioprocessing feature anoxic fermentations that naturally maximize product formation relative to growth of the microbial catalyst. *Vibrio natriegens* is a facultatively fermentative bacterium that has gained attention for bioprocessing due to its rapid growth rate and ease of genetic engineering. However, the fermentative properties of *V. natriegens* have not been compared to traditional bioprocessing workhorses like *Escherichia coli*. We revealed that *V. natriegens* is comparatively sensitive to its own acidic fermentation products, likely because *V. natriegens* lacks acid resistance mechanisms possessed by *E. coli*. Thus, fermentative applications must address this sensitivity either by buffering the fermentations, engineering resistance mechanisms, or bypassing the sensitivity by engineering *V. natriegens* to produce neutral products.

## INTRODUCTION

Bioprocessing utilizes native or engineered microbes to generate value-added products from renewable resources, offering sustainable alternatives to petroleum-based counterparts (1–5). A diverse set of microbes have been proposed as platforms, or chassis, for bioprocessing (6). The marine bacterium *Vibrio natriegens* has increasingly gained attention as a chassis for bioprocessing primarily due to its rapid growth and metabolism (7–9) and the relative ease with which combinatorial genetics can be performed (10–13). Proof-of-concept studies have demonstrated that *V. natriegens* can been engineered to produce a variety of value-added products from a range of carbon substrates (9). However, whereas many bioprocesses use anoxic fermentative conditions to maximize product yields and minimize microbial growth, most *V. natriegens* studies have used oxic conditions, with a few exceptions (8, 14, 15).

Under anoxic conditions with glucose, *V. natriegens* carries out a mixed-acid fermentation, excreting ethanol and a variety of organic acids (Fig 1) (8). The accumulation of organic acids acidifies the medium. In general, acid stress negatively impacts metabolism, viability, and product yield (16). Even under aerobic conditions, *V. natriegens* growth is impacted by acidification from acetic acid that accumulates due to overflow metabolism (17). Thus, it is expected that negative impacts would be exacerbated under anoxic fermentative conditions where there is a greater accumulation of organic acids.

**Fig 1.**
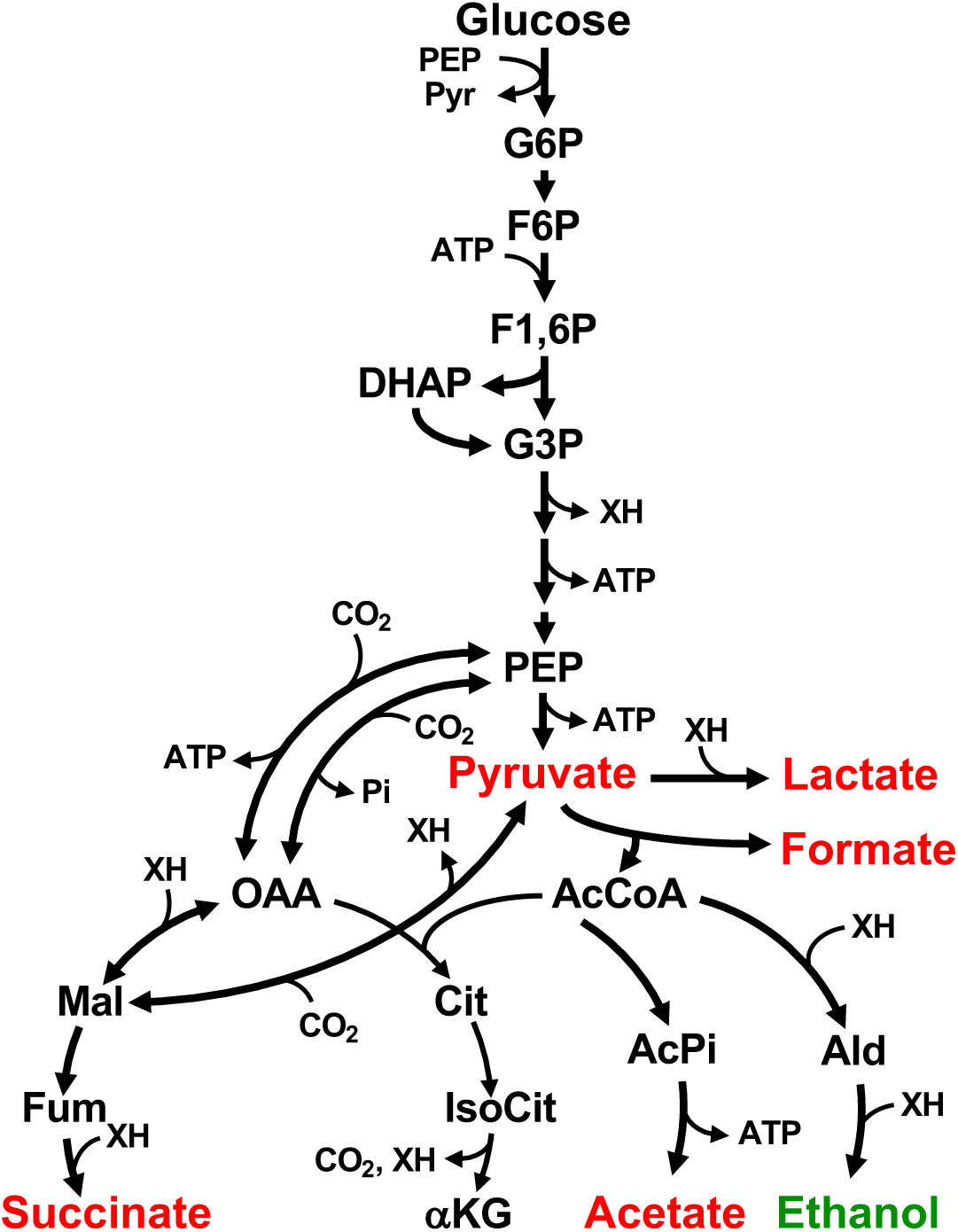
*V. natriegens* fermentative metabolism. Reactions are based on NCBI gene annotations. Red, acidic products (named here for their carboxylate forms); green, neutral products.

Whereas other *Vibrio*s like *V. cholerae* can switch to neutral fermentation products in response to acidic conditions (18), production of neutral fermentation products by *V. natriegens* has not been observed (8). Fermentative bacteria like *Escherichia coli* have an array of mechanisms to deal with acid stress (19, 20). For example, *E. coli* can offset acidosis by converting formic acid into H2 and CO2 using formate hydrogen lyase (FHL), converting an organic acid into two gasses (21). In the presence of certain amino acids, *E. coli* can also eliminate cytoplasmic protons via amino acid decarboxylation reactions (22). As the medium acidifies during fermentation, *E. coli* also shifts its fermentation profile towards lactic acid (23), which might also offset acidosis by producing a single acid instead of the combination of acetic and formic acid (24). The extent to which *V. natriegens* can use these or other acid resistance mechanisms during mixed acid fermentation is unknown.

Here, we compared the growth and metabolic parameters of *V. natriegens* versus *E. coli* under anoxic fermentative conditions. Whereas both organisms produced similar fermentation products, *E. coli* cultures consumed more glucose, accumulated more organic acids, and exhibited longer stationary phase viability. We conclude that *V. natriegens* is comparatively sensitive to fermentative organic acids due to a lack of acid resistance mechanisms.

## RESULTS and DISCUSSION

### *V. natriegens* and *E. coli* have similar fermentative product yields

*V. natriegens* is often compared to *E. coli*, a bacterium that is well-established in synthetic biology, as a potential platform for bioprocessing (7). However, a direct comparison of growth and metabolic parameters under fermentative conditions has not been reported. We grew *E. coli* MG1655 and *V. natriegens* ATCC14048 in their respective anoxic minimal media with 25 mM glucose as the sole carbon source. The media only differed by a higher NaCl concentration for *V. natriegens*, and an optimum growth temperature of 30°C for *V. natriegens* versus 37°C for *E. coli* (Fig 2). Despite the lower growth temperature, *V. natriegens* grew 1.3-times faster than *E. coli* (Fig 2A-C). Although *V. natriegens* reached a higher turbidity (OD660; Fig 2D), *E. coli* has a higher ratio of CFU ml^-1^ : OD660 (Fig 2E) and was thus estimated to achieve a 1.2-fold higher actual cell density (CFU ml^-1^) than *V. natriegens* (Fig 2F).

**Fig 2.**
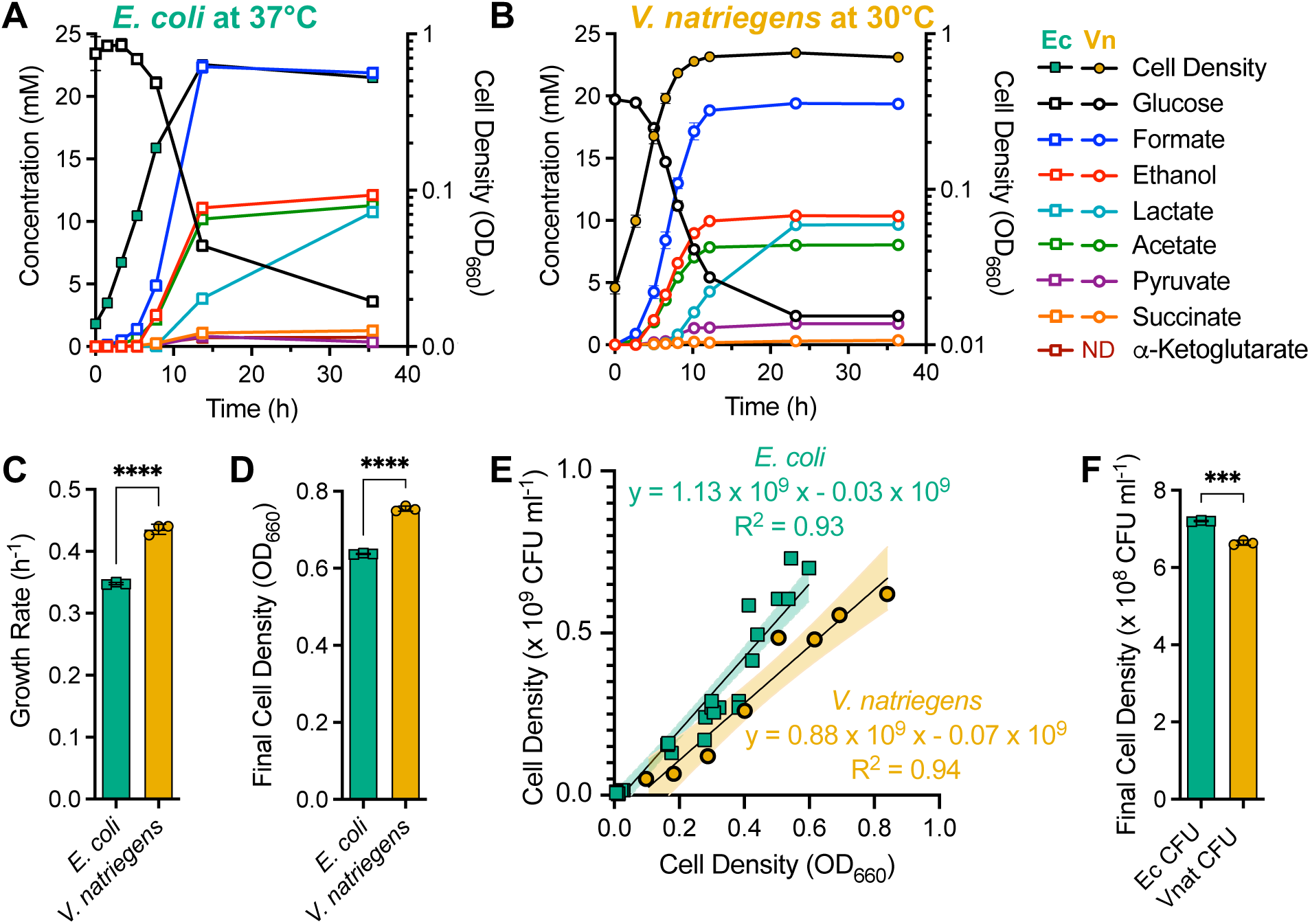
*V. natriegens* grows faster but to a lower cell density while producing a similar array of fermentation products as *E. coli*. *E. coli*. **(A)** and *V. natriegens* **(B)** were grown in 60 ml of anoxic minimal media at their optimum growth temperatures and growth, glucose, and fermentation products were monitored. The same cultures were used to determine exponential growth rates **(C)** and final (highest) cell densities **(D)**. Standard curves to convert turbidity (OD600) measurements of cell density to CFU ml^-1^ **(E)** suggests that the difference in final cell densities between the two species is opposite when considering turbidity **(D)** versus CFU ml^-1^ **(F)**. Error bars, SD; n = 3. Statistical differences were determined using an unpaired, two-tailed t test; ***, *P* < 0.001; ****, *P* < 0.0001.

Late stationary phase (36 h) fermentation product yields between *E. coli* and *V. natriegens* were either statistically similar or exhibited small differences for the most abundant fermentation products: formic acid, no difference; ethanol, no difference; lactic acid, 1.06-fold higher for *V. natriegens*; acetic acid, 1.20-fold higher for *E. coli* (Fig 3A). Both species produced lactic acid as they exited exponential phase, with continued production into stationary phase (Fig 2A-B). Greater fold-differences were observed for minor fermentation product concentrations (Fig 2A, B) and yields (Fig 3A): (i) *E. coli* produced ∼ 1.3 mM succinic acid whereas levels from *V. natriegens* were near the detection limit, (ii) *V. natriegens* produced ∼ 1.7 mM pyruvate that was stable in the supernatant whereas *E. coli* transiently produced as much as 0.8 mM pyruvate, and (iii) *E. coli* produced ∼0.8 mM α-ketoglutarate, whereas none was detected in *V. natriegens* supernatants (Fig 2A-B). Another key difference was that *V. natriegens* did not produce H2 whereas *E. coli* is known to produce H2 as the pH drops below neutral (25, 26). We thus omitted gaseous fermentation products from this study.

**Fig 3.**
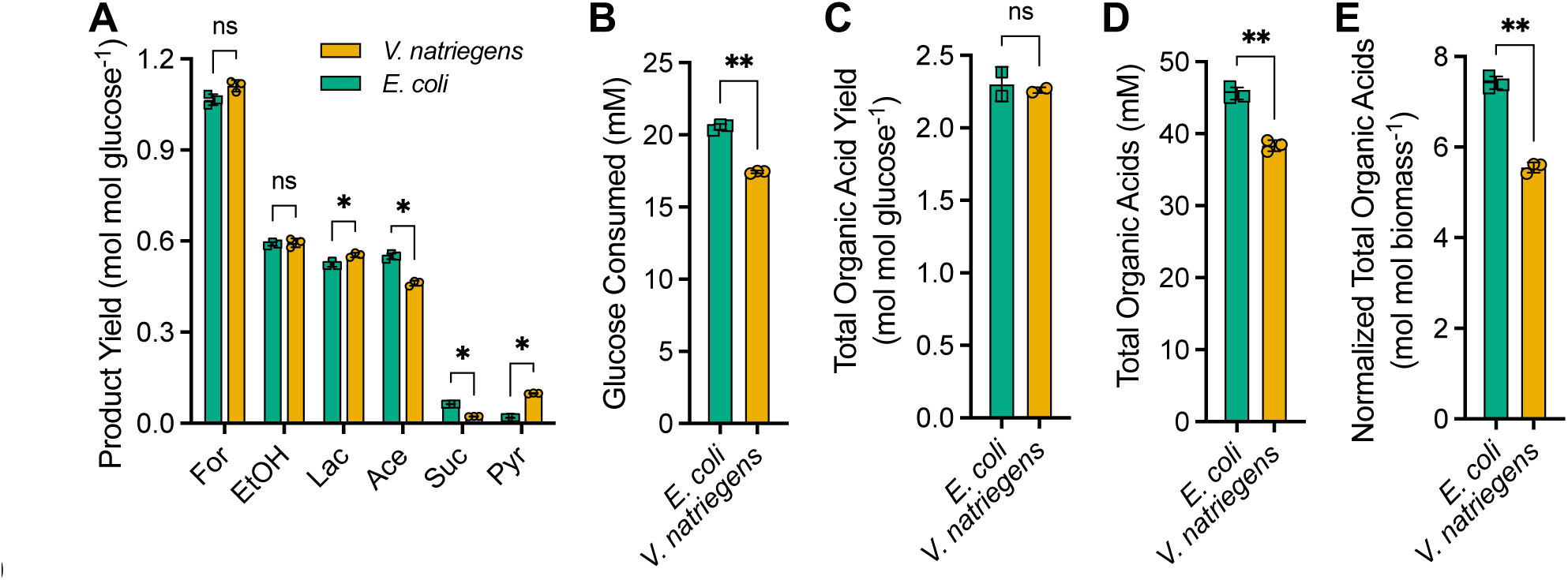
*V. natriegens* consumes less glucose and produces less organic acids than *E. coli*. Fermentation product yields **(A)**, glucose consumed **(B)**, and total organic acid yields **(C)**, concentrations **(D)**, and concentrations normalized for cell density **(E)**, were determined using stationary phase (36 h) samples from Fig 2 cultures. Error bars, SD; n = 3. Statistical differences were determined using an unpaired, two-tailed t test; ns, not significant; *, *P* < 0.05; **, *P* < 0.01. For, formic acid; EtOH, ethanol; Lac, lactic acid; Ace, acetic acid; Suc, succinic acid; Pyr, pyruvate.

### *V. natriegens* is more sensitive to acid than *E. coli*

Neither strain fully consumed the glucose provided (Fig 2A, B) but *E. coli* consumed 1.2-times more glucose than *V. natriegens* (Fig 3B). A possible reason is that each species reached their acid tolerance threshold and *V. natriegens* has a lower acid tolerance. Indeed, although each species had a similar total organic acid yield (Fig 3C), *E. coli* accumulated 1.2-fold more organic acids (Fig 3D) and 1.3-fold more organic acids per cell (Fig 3E).

To compare the acid tolerance between the two species, we tracked cell viability by colony forming units (CFUs) during stationary phase (Fig 4A). Although cell death was not evident from turbidity measurements, CFU measurements indicated that both species ultimately went extinct. However, whereas viable *E. coli* cells were detected up to 82 h after inoculation, no viable *V. natriegens* cells were detected 24 h after inoculation, about 12 h after the end of the exponential growth phase (Fig 4A).

**Fig 4.**
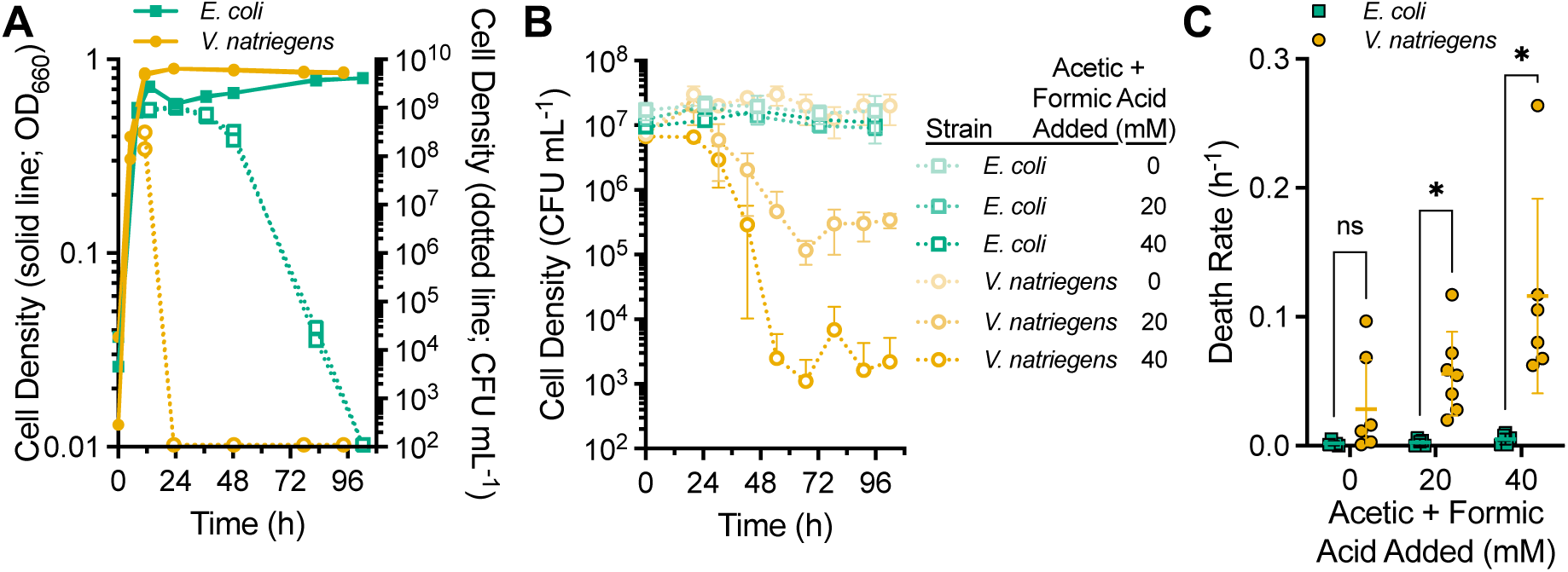
*V. natriegens* is sensitive to acidic fermentation products compared to *E. coli*. **(A)** Culture turbidity (solid lines; OD660) was tracked in minimal media with 25 mM glucose over the entire experiment whereas viable cell density (dotted lines; CFU ml^-1^) was tracked from the end of the growth phase (∼12 h). All data points from the duplicate cultures are shown. **(B)** Viable cell counts were monitored in cell suspensions inoculated in late exponential phase to defined media lacking glucose with different concentrations of a 1:1 solution of acetic and formic acid. Data points, mean ± SD; n=3. (**B,C**) The limit of detection is 10^2^ CFU ml^-1^. **(C)** Death rates determined from the slope of a semi-log plot fitted to the data from panel B and two other similar experiments using different starting cell densities and incubation times. Horizonal line, mean ± SD; n=5-7. Statistical differences were determined using unpaired, two-tailed t tests; ns, not significant; *, *P* < 0.05.

To verify that organic acids were a primary driver of *V. natriegens* cell death, we inoculated late exponential phase cultures (*V. natriegens*, 0.65-0.69 OD660 ; *E. coli*, 0.82-0.87 OD660) to media lacking glucose but with different concentrations of a 1:1 mixture of acetic and formic acid. The cultures used to prepare these suspensions were grown with less glucose (15 mM) to ensure less acid exposure prior to the assay. Without added organic acids, both species showed little decline in cell viability (Fig 4B, C). *E. coli* viability was also unaffected by 20 and 40 mM organic acids (Fig 4B, C). In contrast, increasing the organic acid concentration progressively decreased *V. natriegens* viability (Fig 4B) and increased the death rate (Fig 4C).

We then verified that the *V. natriegens* sensitivity to organic acids is dependent on the pH by increasing the buffering capacity of medium with either 100 or 200 mM 4-morpholinepropanesulfonic acid (MOPS) buffer. In each case, viable cell counts exceeded 10^8^ CFU mL^-1^ at the onset of stationary phase, even though the turbidity was notably lower with 200 mM MOPS compared to the other conditions (Fig 5A). Unlike the rapid extinction observed when MOPS was excluded, 100 mM MOPS delayed extinction by 24 h and 200 mM MOPS avoided extinction within the 100-h time course (Fig 5A). Even though cells still went extinct with 100 mM MOPS, glucose was fully consumed by 24 h, with continued production of most fermentation products, especially lactic acid (Fig 5B, C). Thus, we conclude that the rapid extinction of *V. natriegens* upon reaching stationary phase under fermentative cultures is due to acid sensitivity.

**Fig 5.**
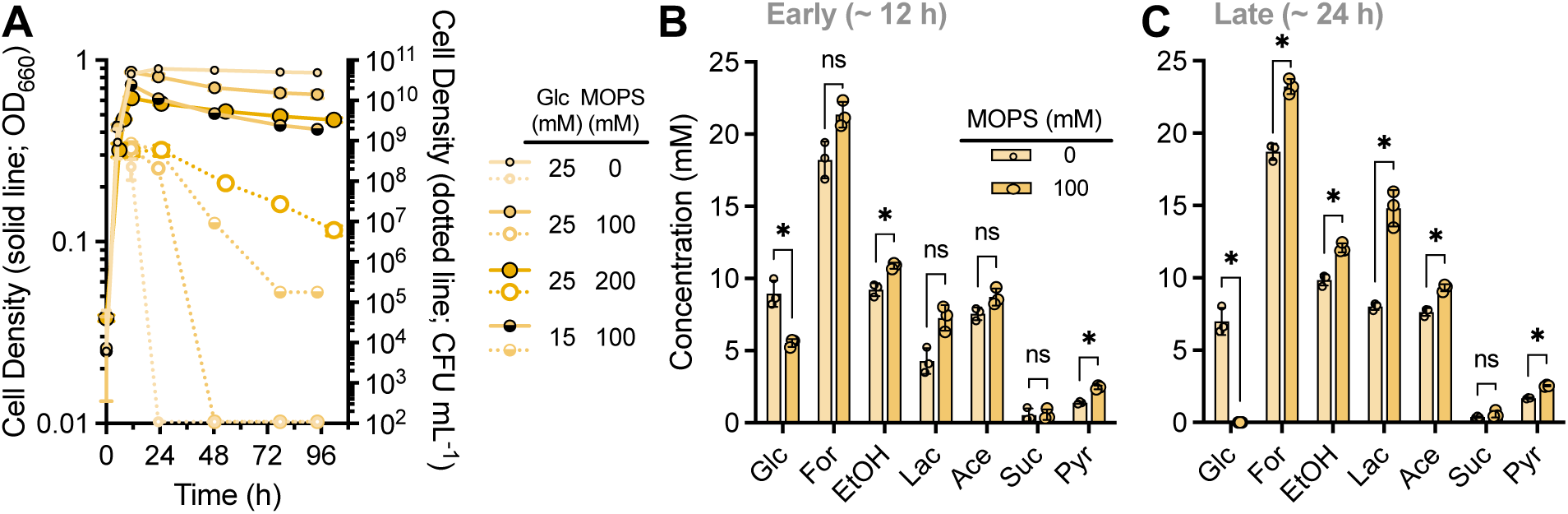
Increasing the buffer concentration of the growth medium alleviates *V. natriegens* organic acid sensitivity. **(A)** *V. natriegens* culture turbidity (solid lines; OD660) was tracked in minimal media with different concentrations of glucose and MOPS over the entire experiment whereas viable cell density (dotted lines; CFU ml^-1^) was tracked from the end of the growth phase (∼12 h). The ‘0 mM MOPS’ data is the same as in Fig 4A. Each data point represents the mean ± range; n=2. The limit of detection is 10^2^ CFU ml^-1^. **(B,C)** Remaining glucose and accumulated fermentation products from *V. natriegens* cultures grown with 25 mM glucose in ‘early’ stationary phase (**B**, ∼ 12 h) and ‘late’ stationary phase (**c**, ∼ 24 h). Error bars, mean ± SD; n=3. Statistical differences were determined using unpaired, two-tailed t tests; ns, not significant; *, *P* < 0.05. Glc, glucose; For, formic acid; EtOH, ethanol; Lac, lactic acid; Ace, acetic acid; Suc, succinic acid; Pyr, pyruvate.

### *V. natriegens* lacks acid resistance mechanisms

We sought an explanation for the stark difference between *V. natriegens* and *E. coli* survival in acidic conditions. Bacteria like *E. coli* and *V. cholerae* have mechanisms to deal with acid stress both during fermentative conditions and to transit the highly acidic stomach (19, 20). To determine the acid resistance repertoire of *V. natriegens*, we performed BLASTp searches (27) using *E. coli* and *V. cholerae* acid resistance proteins as query sequences (Table 1).

**Table 1.**
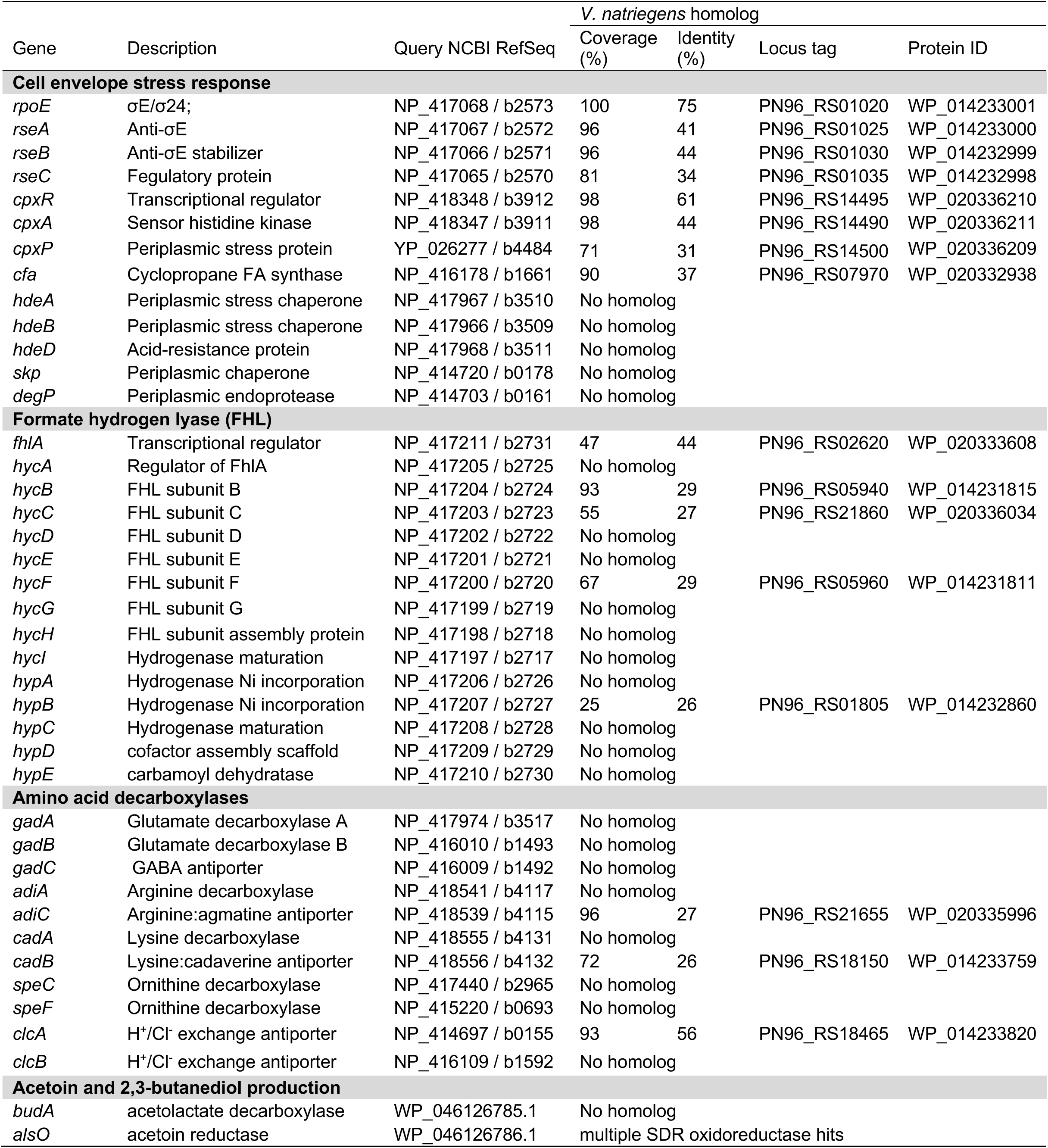
*V. natriegens* acid tolerance proteins.

Based on our homolog threshold criteria of 25% identity over 50% of the query sequence, *V. natriegens* likely has several proteins to respond to cell envelope stress, which could include acid stress, but also other stressors, like heat. For example, we identified a σ^E^ homolog and associated proteins that, in *E. coli*, activates a regulon in response to denatured outer membrane proteins, and the Cpx two component system that responds to periplasmic and inner membrane stress signals (28). Among genes of the *E. coli* Cpx regulon, *V. natriegens* also has *cfa*, which encodes a protein that produces cyclopropane fatty acids that can protect against various stressors, including acidity (29). However, *V. natriegens* appears to lack many of the chaperones and proteases (i.e., *degP*, *hdeABD*, *skp*) that are crucial in *E. coli* for the proper folding or degradation of cell envelope proteins under stress conditions (30–32).

When considering specific responses to acid stress, the *V. natriegens* genome suggests that it is even less well-equipped. FHL, a complex of formate dehydrogenase (FDH) and hydrogenase, converts formic acid into CO2 and H2, thereby converting an acidic fermentation product into two gasses (21, 25). Whereas *V. natriegens* has FDH protein homologs, *V. natriegens* does not have hydrogenase genes. In agreement with this bioinformatic result, no H2 was detected in *V. natriegens* fermentative cultures by gas chromatography. *V. natriegens* also lacks amino acid decarboxylases that can eliminate cytoplasmic protons (19, 20, 22), but it has one of two H^+^/Cl^-^ antiporters that help prevent hyperpolarization of the membrane during the use of amino acid decarboxylases (33). Regardless, we would not expect amino acid decarboxylases to be effective in our minimal medium that did not contain amino acids.

### Lactic acid production prolongs growth and metabolism but does not improve survival

Organisms that produce multiple fermentation products can alter their fermentation profile to respond to environmental conditions. Unlike *V. cholerae*, *V. natriegens* lacks genes for producing neutral fermentation products (Table 1) (18). However, like *E. coli*, *V. natriegens* produced lactic acid starting in late exponential phase and into stationary phase (Fig 2A-B). In *E. coli*, cytoplasmic lactate dehydrogenase (LDH) expression increases in response to acidic pH (23), suggesting that lactic acid production might decrease acid stress (24). Shifting to lactic acid production would result in less ATP but also less acid overall to maintain electron balance (Fig 6A); 2 pyruvate could either go to 2 lactatic acid (pKa, 3.86) or to 1 ethanol + 2 formic acid (pKa, 3.75) + 1 acetate (pKa, 4.76). The latter pathway would generate more organic acids, one of which is acetic acid that would more readily deprotonate due to its high pKa, and which is potentially more toxic than lactic acid in general (34).

**Fig 6.**
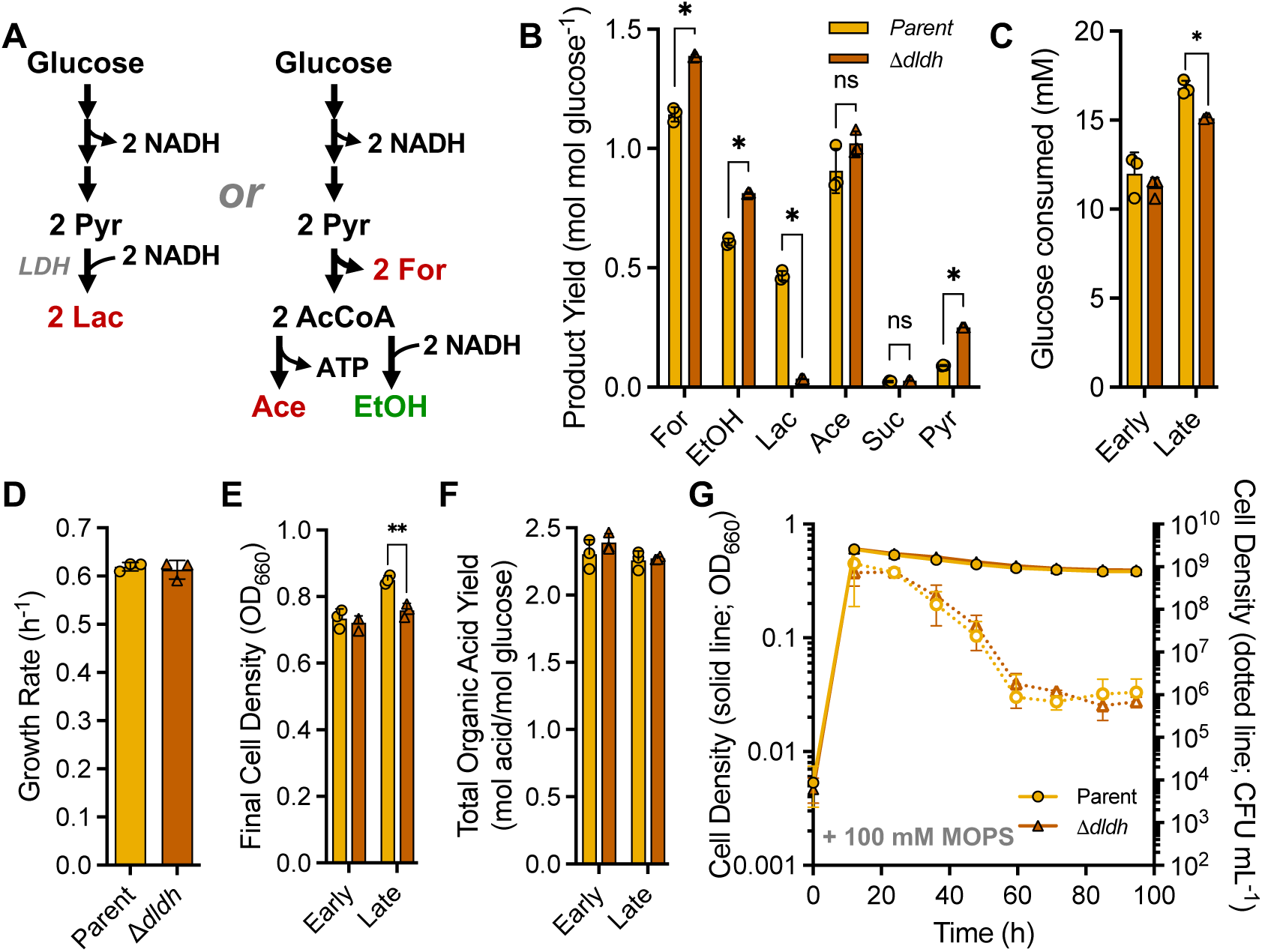
*V. natriegens* LDH extends fermentative metabolism and growth but not viability. **(A)** Lactic acid production can hypothetically achieve electron balance while producing less acid (2 lactic acid; Lac; left) via LDH compared to producing ethanol, formic acid, and acetic acid (right). Comparison of ‘late’ (24 h) fermentation product yields **(B)**, glucose consumed **(C)**, exponential growth rate **(D)**, final (highest) cell density **(E)**, and total organic acid yield **(F)** between the parent strain (WT Δ*dns*::Spec^R^) and the LDH mutant (Δ*dldh*) in a minimal medium with 25 mM glucose and no MOPS. For, formic acid; EtOH, ethanol; Lac, lactic acid; Ace, acetic acid; Suc, succinic acid; Pyr, pyruvate. **(C,E,F)** Early, 9 h exponential phase samples; Late, 24 h stationary phase samples. **(G)** Turbidity (solid lines; OD660) and viable cell density (dotted lines; CFU ml^-1^) were tracked into stationary phase in minimal medium with 25 mM glucose plus 100 mM MOPS; each data point is the mean ± SD. The limit of detection is 10^2^ CFU ml^-1^. **(B-G)** Error bars, SD; n=3. Statistical differences were determined using unpaired, two-tailed t tests; ns, not significant; *, *P* < 0.05; **, *P* < 0.01.

To test whether lactic acid production contributes to *V. natriegens* survival under fermentative conditions, we first created an LDH mutant. *V. natriegens* has three different putative LDH genes, *BA890_15740*, *BA890_20630* (*dldh*), and *BA890_20645* (*lldh*). The latter two genes had been deleted by another group and had little effect on fermentative lactic acid production (8). However, the conditions in that study favored succinic acid production over lactic acid production whereas we observed the opposite. Using the fermentative LDH gene from *E. coli* MG1655 (P52643) as a BLASTp query, we hypothesized that *dldh* was most likely the *V. natriegens* fermentative LDH (100% coverage, 62% identity); the other two candidates did not show significant sequence identity to the query sequence. Indeed, when we deleted *V. natriegens dldh*, the lactic acid yield was close to zero at 24 h (Fig 6B). Compared to the parent strain, the Δ*dldh* mutant produced 1.2-times more formic acid and 1.3-times more ethanol (Fig 6B), a route that would replace the electron balancing role of lactic acid production (Fig 6A). The Δ*dldh* mutant also produced 2.8-times more pyruvate, suggesting that a bottleneck was created in the absence of LDH. The Δ*dldh* mutant consumed less glucose than the parent (Fig 6C), which did not affect the growth rate (Fig 6D), but the final Δ*dldh* mutant turbidity was 90% of that of the parent (Fig 6E). Still, the total organic acid yield was not significantly different between the two strains, suggesting that even without lactic acid production, *V. natriegens* faced comparable acid exposure (Fig 6F).

We thus examined the effect of the LDH deletion on *V. natriegens* stationary phase survival under fermentative conditions. Anticipating that the rapid death rate would make a comparative analysis difficult, we added 100 mM MOPS to slow the death rate; MOPS does not affect stationary phase lactic acid production (Fig 5B, C). Stationary-phase survival was similar between the parent and Δ*dldh* strains (Fig 6G). Thus, whereas lactic acid production improves growth (Fig. 6E, G) and extends glucose consumption into stationary phase, it does not contribute to acid tolerance under fermentative conditions.

Below we speculate why lactic acid might be produced during the transition out of exponential growth and into stationary phase. One possibility is that acetate and ethanol production become less available as an option. For example, as the pH drops and growth slows, the biosynthetic demand for ATP would also slow. This might limit ADP availability for acetate production and create a bottleneck at pyruvate (Fig 6A). LDH offers a carbon and electron sink to maintain metabolic flow and help alleviate this bottleneck (Fig 6A); a role for LDH in addressing this bottleneck would also explain the elevated pyruvate excretion in the Δ*dldh* mutant (Fig 6B). If we assume similar kinetic parameters to *E. coli* enzymes, LDH is poised to respond to pyruvate accumulation and low pH; as the pH drops from neutral to 5, LDH activity increases 1.6-fold and the relatively low affinity (Km) for pyruvate improves from 7.2 to 5.0 mM. Although PFL has a higher affinity for pyruvate (Km, 2.0 mM) the optimal pH for activity is 7.5, and would thus worsen if the cytoplasm begins to acidify (35–37).

## Conclusion

As interest grows in *V. natriegens* as a bioprocessing strain, one must weigh its favorable attributes of rapid growth and genetic tractability against its less favorable attributes. Here we report that, relative to *E. coli*, *V. natriegens* is comparatively sensitive to the acidic conditions that arise during mixed acid fermentations. This difference in acid tolerance is likely due to *V. natriegens* lacking most of the acid resistance mechanisms that have been characterized in *E. coli*. Such acid-resistance mechanisms could potentially be engineered to improve *V. natriegens* acid tolerance. Increasing the buffering capacity of the fermentation medium would also avert acid sensitivity, but one would need to consider the cost of buffer under industrial conditions. Thus, acid sensitivity will need to be considered when engineering *V. natriegens* for bioprocessing, unless the goal is to engineer *V. natriegens* to produce neutral products.

## MATERIALS AND METHODS

### Strains

Experiments used the wild-type strains *Escherichia coli* MG1655 (38) and *Vibrio natriegens* ATCC 14048 (10) or its derivatives. *V. natriegens* mutants were made as described (10), except the transformation mixtures of cells and PCR products were incubated statically at 30°C for 12-16 h before outgrowth, and selection plates used 250 µg mL^-1^ spectinomycin. The *V. natriegens* Δ*dldh* mutant was made in *V. natriegens* TND1964, which is the wild-type strain with pMMB-*tfoX* that allows for IPTG-inducible competency (10). Briefly, a Δ*dldh* (Δ20530) construct was made by PCR amplifying (Q5 DNA polymerase; New England Biolabs) 3kb upstream and downstream fragments. The two fragments were then combined by splicing-by-overlap extension PCR using the outermost primers. The Δ*dldh* construct (1 µg) was then co-transformed into TND1964 with 50 ng of a DNA fragment containing a spectinomycin resistance cassette flanked by ∼1 kb of homology to the upstream and downstream regions of the *dns* locus (*BA890_12415*)amplified from *V. natriegens* SAD1306 genomic DNA (10). The ‘parent’ strain (NWH004) used as the control strain in experiments with the Δ*dldh* mutant was derived from TND1964 by introducing a spectinomycin resistance cassette at the *dns* locus. Colonies were screened by PCR using OneTaq DNA polymerase (New England Biolabs). Mutations were confirmed by Sanger sequencing. Primers are in Table 2.

**Table 2.**
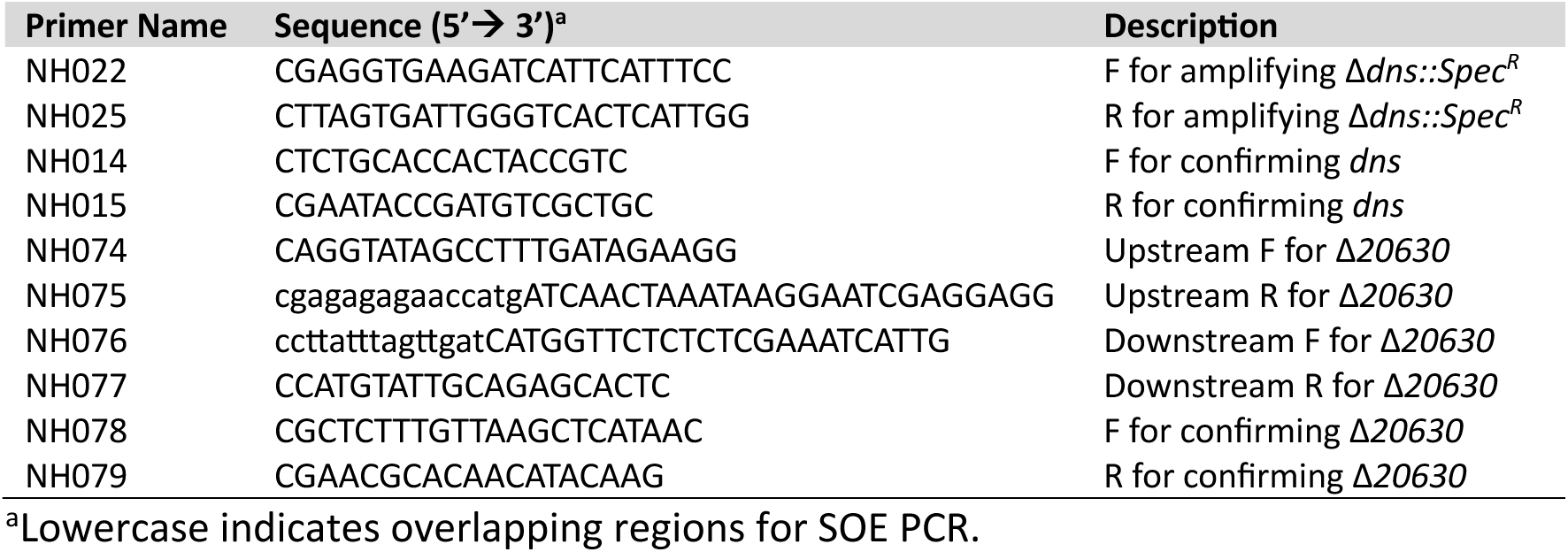
Primers used in this study.

### Growth conditions

Colonies were grown from 25% glycerol frozen stocks on agar plates of either lysogeny broth medium (LB; *E. coli*), or the media supplemented with an additional 20 g NaCl L^-1^ (LB3; *V. natriegens*). LB3 was supplemented with either 100 µg mL^-1^ carbenicillin, 250 µg mL^-1^ spectinomycin, or 100 µM IPTG when appropriate. LB3 was also used routinely for strain construction.

For all other experiments anoxic minimal M9-derived coculture medium (MDC) (39) for *E. coli* or MDC with an additional 80 mM NaCl for *V. natriegens* (MDC80) was prepared by aliquoting 10 ml media into 27-ml anaerobic tubes or 60 ml media into 150-ml serum vials, bubbling with N2, then sealing with rubber stoppers and aluminum crimps before autoclaving. Media supplements were then added by syringe to the following final concentrations from anoxic 100X stock solutions: glucose, 25 mM; NH4Cl, 10mM; cation solution 1% v/v (1 mM MgSO4 and 0.1 mM CaCl2). A 2 M anoxic MOPS stock solution (pH 7.0) was filter sterilized into a serum vial and then flushed with filter-sterilized N2 and then added to cultures to the indicated concentration after autoclaving, with sterile ultrapure water also added where necessary to have common dilution effects between cultures. Starter cultures were inoculated to MDC or MDC80 from single colonies. Late-exponential phase starter cultures (0.65-0.85 OD660) were then used to inoculate test media to an initial cell density of ∼0.01 OD660. Tubes were incubated horizontally at either 30°C (*V. natriegens*) or 37°C (*E. coli*) with shaking at 150 rpm.

### Analytical procedures

A Genesys 20 spectrophotometer (Thermo-Fisher) was used to measure cell density at 660 nm (OD660) either directly in the 27-ml culture tubes or in 1-ml samples in cuvettes when 150-ml serum vials were used. Growth rates were determined by fitting an exponential trendline to turbidity measurements taken between 0.1-0.8 OD660 using GraphPad Prism v6.

Glucose and fermentation products were quantified in supernatant samples by HPLC (Shimadzu) as described (40). Temporal assessment of fermentations was performed in 60-mL cultures in 150-mL serum vials. Samples (1 ml) were taken at regular intervals; air contamination was minimized by flushing a 1-mL syringe and needle with filter-sterilized N2 before sampling. To check for H2, 0.1 ml of headspace gas was sampled using a gas-tight syringe and injected into a Shimadzu GC-2014 gas chromatograph with a thermal conductivity detector as described (41).

Viable cell counts were quantified as CFUs using the track plate method (42). Samples (0.4-0.5 ml) were then removed from 10-ml cultures using an N2-flushed syringe and serially diluted in oxic MDC or MDC80 in a 96-well plate. The last six dilutions were spotted (10 µL) onto a LB or LB3 agar using a multichannel pipette. A minimum of 1 colony a cutoff, making the limit of detection 100 CFU mL^-1^ for an undiluted sample.

For experiments using nongrowing cells, late-exponential phase cultures (0.65-0.85 OD660) were inoculated to anoxic MDC or MDC80 with all supplements except glucose. The target inoculum was 10^7^ cells mL^-1^. The pH was decreased by adding the appropriate volume of a solution containing 1 M glacial acetic acid plus 1 M formic acid by syringe. Cell suspensions were then sampled for CFU measurements. Death rates were determined from the slope of a semi-log equation (linear X, log Y) fitted to experimental data using GraphPad Prism v6.

### Bioinformatics

NCBI’s BLASTp (27) was used to identify putative acid-resistance genes in *V. natriegens* ATCC14048 (NCBI RefSeq: GCF_001456255.1) using amino acid sequences from *E. coli* MG1655 (NCBI RefSeq: GCF_000005845.2) or *V. cholerae* (GCF_000969265.1). *V. natriegens* proteins that had >25% identity with >50% query coverage were considered putative homologs. **Statistics.** GraphPad Prism v6 was used for all statistical analyses.

## Acknowledgments

This work was supported in part by the National Science Foundation CAREER award MCB-1749489. NWH was supported by a IU College of Arts and Sciences John and Wendy Kindig Fellowship. We thank A. Dalia and the Dalia lab for bacterial strains, resources, and guidance for *V. natriegens* genetics. We also thank C. Fuqua, C. Landeta, and the McKinlay lab for helpful discussions.

We declare no conflicts of interest.

